# Nectophore coordination and kinematics by physonect siphonophores

**DOI:** 10.1101/2023.04.12.536580

**Authors:** Shirah Strock, John H. Costello, Joost Daniels, Kakani Katija, Sean P. Colin

## Abstract

Siphonophores are ubiquitous and often highly abundant members of pelagic ecosystems throughout the open ocean. They are unique among animal taxa in that they use multiple jets for propulsion. Little is known about kinematics of the individual jets produced by nectophores or how the jets are coordinated during normal swimming behavior. Using remotely operated vehicles and SCUBA, we video recorded the swimming behavior of several physonect species in their natural environment. The pulsed kinematics of the individual nectophores that comprise the siphonophore nectosome were quantified and, based on these kinematics, we examined the coordination of adjacent nectophores. We found for all species that nectophores sharing the same side of the nectosomal axis were coordinated metachronally. However, this coordination was not shared with nectophores on the opposite side of the nectosomal axis. For most species, the metachronal contraction waves of nectophores were initiated by the apical nectophores and traveled dorsally. However, the metachronal wave of *Apolemia rubriversa* traveled in the opposite direction. Although nectophore groups on opposite sides of the nectosome were not coordinated, they pulsed with similar frequencies. This enabled siphonophores to maintain relatively linear trajectories during swimming. The timing and characteristics of the metachronal coordination of pulsed jets affects how the jet wakes interact and may provide important insight into how interacting jets may be optimized for efficient propulsion.

## Introduction

Physonect siphonophores are colonial cnidarians that are found throughout the pelagic communities of the open ocean. In these communities, siphonophores are often dominant predators that have important trophic roles and impacts (Pugh, 1975; Purcell, 1981). For example, in Monterey Bay, California, *Nanomia bijuga,* a physonect siphonophore, is often the most commonly observed gelatinous animal throughout the year (Robison et al., 1998). They eat crustacean zooplankton, and in the Monterey Bay pelagic communities they are important predators of *Euphausia pacifica, E. hysanoessa* spp., and *Nematoscelis difficilis* (Robison et al., 1998). Most siphonophores forage as ambush predators and spend most of their time drifting motionlessly with their tentacles deployed. For these siphonophores, swimming by the nectosome is used for repositioning, migration and predator escape. However, in situ observations also suggest that some species may swim more continuously, pulling deployed tentacles behind the nectosome. The different uses of swimming, times spent swimming and ability of the nectosome to transport the siphosome are all factors that will impact the trophic role and impact of different taxa (Purcell, 1980).

Siphonophores are unique in the animal kingdom because they are one of two taxa that utilize multiple jets for swimming propulsion (Costello et al., 2015). These jets are produced by propulsive colony members known as nectophores (Figure 1B), which use muscular contractions to create jets of water that propel the whole organism through the water column. The nectophores are genetically identical clones and are arranged to form a coherent unit called the nectosome (Figure 1B). During whole colony swimming, the nectosome pulls the feeding and reproductive colony members that comprise the siphosome (Figure 1B). While the nectosome propels the colony through the water, forward propulsion of the siphonophore is fundamentally driven by individual-scale contractions of the nectophores (Sutherland et al., 2019a). Understanding the individual motions of the nectophores is required for understanding the swimming kinematics of the entire colony.

**Fig 1.**
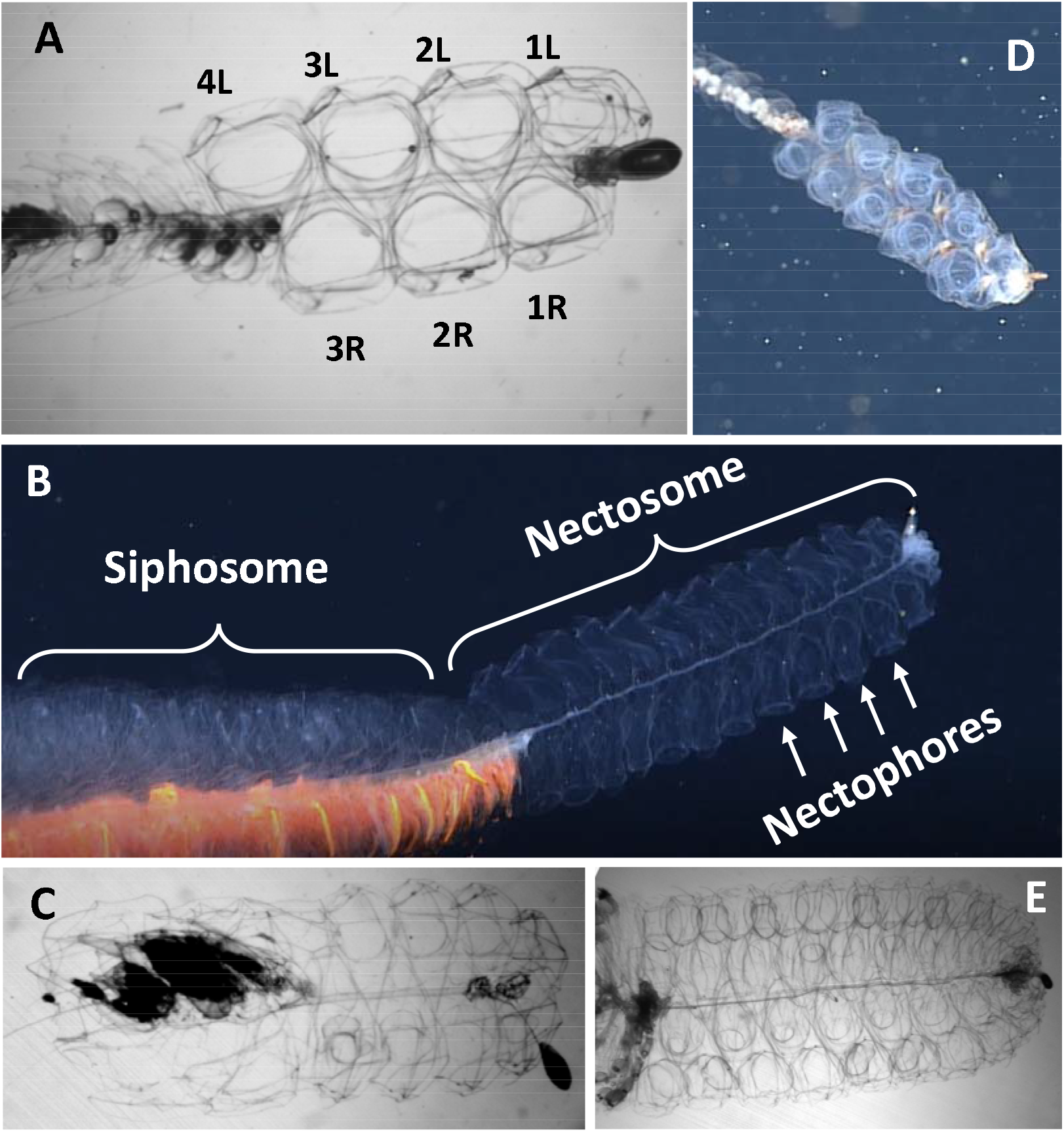
Morphology of physonect siphonophores. A) Image of *Nanomia bijuga* illustrating how individual nectophores were labeled for analysis. B) Images of the Galaxy siphonophore illustrating different parts of the siphonophore colony. C, D and E) *Algama okeni, Apolemia rubriversa* and Forskalia sp. swimming in the field, respectively.

Physonectae is a suborder of siphonophores that are highly diverse morphologically, functionally and ecologically. Both the number and morphology of the nectophores vary greatly among physonects, and these differences impact the swimming proficiency and efficiency of the nectosome (Du Clos et al., 2022; Jiang et al., 2021) and, by extension, how well the nectosomes function for different behaviors. However, no data exists on the relationships of morphology with swimming abilities or behavioral strategies of siphonophores. This is primarily due to the limited number of species that can be effectively captured for analysis, but also due to the high diversity in morphology among the different species of siphonophores. Furthermore, the complexities of the multi-jet propulsion performed by these organisms poses a challenge in terms of understanding the function of these species. Understanding how these siphonophores use their nectophores to swim is the first step in developing a functional understanding of their diverse ecological roles and impacts.

The goal of this study was to investigate how morphologically distinct physonect siphonophores coordinate their jetting nectophores for normal, non-escape swimming. This was done using in situ video recordings of different species during routine swimming -- non-escape swimming in the forward direction such that the siphosome is trailing behind the nectosome and moving in a straight line. Nectophore kinematics were quantified and compared for 5 physonect species: Galaxy siphonophore (not yet described species), *Apolemia rubriversa, Nanomia* spp. (2 locations), *Forakalia* sp. and *Agalma okeni*.

## Methods

Nectophore kinematics of the physonect species were quantified from videos during “normal” swimming. Most of the data were collected from animals swimming in the field at 2 locations (Monterey Bay, CA, USA and Friday Harbor, WA, USA); one was from the laboratory at a third location (Santa Catalina, Panama, Table 1). Field observations were collected using SCUBA or a remotely operated vehicle (ROV). ROV videos were captured using the ROV *MiniROV* in the Monterey Bay National Marine Sanctuary near Midwater Station 1 (latitude:36° 41.8792 N, longitude: 122° 2.9929 W), with bottom depths exceeding 400 m. Multiple dives with the ROV were made in the spring and summer of 2019 and 2020, and in the autumn of 2019. *MiniROV* is a small, 1500 m-rated electric ROV platform, designed to minimize acoustic and hydrodynamic disturbances so as to minimize organismal disruptions and behavior during observations. The vehicle is equipped with a main science camera (Insite Pacific Incorporated Mini Zeus II), a stereo imaging system (Katija et al. 2021; Allied Vision G-319B monochrome cameras and Marine Imaging Technologies underwater housings with domed-glass optical ports), a pair of red lights (Deep Sea Power and Light MultiRay LED Sealite 2025 at 650–670 nm), and additional vehicle sensors. Red illumination was used whenever possible to minimize disruptions and changes in animal behavior (e.g., avoidance, attraction, escape).

**Table 1.** Summary of kinematic data analyzed for this study, with data on whether the analysis is from a field or lab video, the total number of sequences that were analyzed for each species, and the number of individuals that were analyzed for each species.

| Species | Field or Laboratory/Field method | Number of Individuals Analyzed | Number of Sequences |
| --- | --- | --- | --- |
| Galaxy Siphonophore | Field/ROV | 1 | 7 |
| *Apolemia rubriversa* | Field/ROV | 9 | 10 |
| *Nanomia bijuga* (Friday Harbor) | Field/SCUBA | 5 | 5 |
| *Nanomia* sp. (Panama) | Laboratory | 10 | 10 |
| *Forakalia* sp. | Field/ROV | 10 | 10 |
| *Agalma okeni* | Field/ROV | 5 | 5 |

Nectophore kinematics videos of *Nanomia bijuga* were collected in Friday Harbor by SCUBA divers using a FASTEC TS5 (250 fps, 2560×2046 pixels) and brightfield illumination (Costello et al., 2015). Laboratory videos of *Nanomia* sp. from Panama were taken of individuals freshly hand collected by SCUBA and placed into a gently circulating kreisel tank. Individuals in the tank appeared to have normal field behavior, switching between resting with tentacles deployed and swimming to reposition. The swimming to reposition was the behavior that we defined as routine swimming for subsequent kinematics analysis -- non-escape swimming in the forward direction such that the siphosome is trailing behind the nectosome and moving in a straight line.

The videos were analyzed for multiple variables based on nectophore jetting (Table 1). Nectophores were numbered so they could be differentiated and labeled consistently throughout the study (Fig. 1A). For species where the nectophores are arranged bilaterally along a plane we sequentially labeled the nectophores along each side of the nectosome from the apex to the posterior end. The nectophores were also labeled for whether they appeared in the video to be arranged along the left or right side of the nectosome. The videos were observed frame by frame, and the contraction and relaxation of each individual nectophore was quantified through time. Our goal was to analyze at least 10 swim cycles for each replicate individual, and ≥ 5 replicate individuals were analyzed per species (Table 1). The galaxy siphonophore, a very rarely observed undescribed species, was the exception, and pseudo-replication (n = 7) via analyzed sequences several minutes apart were obtained from a single 10 minute recording.

Several kinematic parameters were quantified including jet frequency (jets/s), interpulse duration (s), and contraction duration (s). Jetting frequencies were quantified as the number of jets per second each nectophore performed. The interpulse duration (or duration time between pulses) was the amount of time in seconds that the nectophore was relaxed and not jetting, or the time between the sequential jets of one nectophore. The pulse duration was the time in seconds that each nectophore was contracted. In addition to individual nectophore kinematics, the timing of contractions of adjacent nectophores or offset time was analyzed. Offset was defined as the amount of time between jets of adjacent consecutive nectophores along a side of the siphonophore, which was determined by subtracting the jet start time of each nectophore from the start of the nectophore contraction immediately behind it on the nectosome.

Overall, there were four analyses completed for each video: jetting frequency, interpulse duration, pulse duration, and offset time. The values for each of these four characteristics were averaged together for all of the individuals of one species and for the left and right nectophores (for all five species, it was found that there was no significant difference between the right and left sides; t-test, p > 0.05, p = 0.0566-0.9923).

Jetting kinematic parameters were compared among species using single factor ANOVAs. All the data conformed to the assumption of normality, however in some cases the data was square root transformed to achieve equal variances. The Tukey Kramer method was used for post-hoc comparisons. omparisons of kinematics between nectophores on either side of the nectosome used T-tests. If these kinematics were not significantly different, then all nectophores on a nectosome were pooled for comparisons among species.

To examine the hydrodynamics of adjacent, asynchronous jets, we used laser sheet particle image velocimetry (PIV) on *N. bijuga* from Friday Harbor, WA. Individual siphonophores were hand collected from the docks surrounding the Friday Harbor laboratories, and immediately transported to the laboratory for PIV videography (Sutherland et al., 2019b). Individuals were placed into glass filming vessels with filtered seawater that was seeded with 10 mm hollow glass beads. The vessel was illuminated with <1 mm thick laser sheet (532 nm) and *N. bijuga* was recorded swimming in the laser sheet at 6400 fps (1024×1024 pixels) using a Photron AX200. It is rare for *N. bijuga* to swim asynchronously when it is in the PIV filming vessel, however, we did record one sequence where an individual swam asynchronously from rest. In this sequence, the plane of the laser passed through the middle of the nectophore and the jet wake. Image pairs of this sequence were analyzed using a cross-correlation PIV algorithm with multi-pass interrogation windows of decreasing size (64 to 32 pixels) and 50% overlap (LaVision DaVis 8.3). At 6400 fps there was no streaking in the jet wake.

## Results

During swimming, all the nectophores along the nectosome contract and relax with repeated cycles over time (Fig. 2A and B). Fig. 2 shows the raw cycle data for the Galaxy siphonophore (Fig. 2A-C) and for *Apolemia rubriversa* (Fig. 2D-E). The raw data illustrates how each nectophore continues its own contraction-relaxation cycle over time. In addition, it shows that nectophores contract and relax asynchronously and, qualitatively, nectophore contraction-relaxation cycles appear to be coordinated at some parts of the nectosome (seen as regular patterns in the raw data). Interestingly, for the Galaxy siphonophore, the anterior 4 nectophores appear less coordinated than the posterior 6 nectophores (Fig. 2B vs C). Nectophore development occurs at the apex (e.g., apical nectophores are less mature than dorsal), and this developmental pattern may affect their ability to coordinate between adjacent nectophores. In addition, apical nectophores have been shown to be primarily used for maneuvering in *N. bijuga* and may not coordinate for steady-state straight swimming (Costello et al., 2015).

**Fig. 2.**
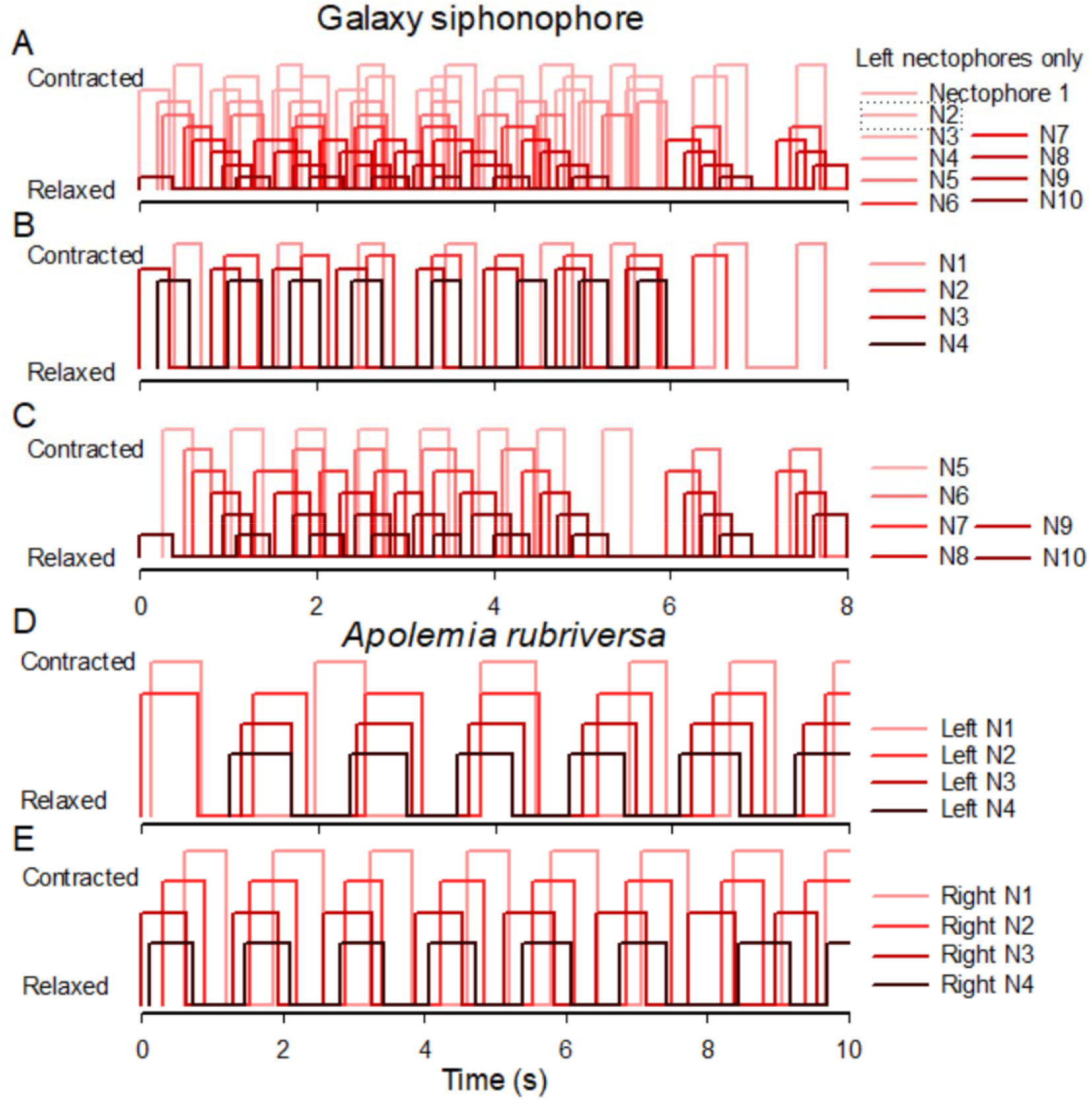
Raw nectophore pulsing data showing the contraction and relaxation of the nectophores of the Galaxy siphonophore (A-C) and *Apolemia rubriversa* (D-E). Nectophores are numbered sequentially starting at the apex of the nectosome. Only showing data for one side of the Galaxy siphonophore.

Kinematic parameters across all five species reveal that some parameters are much less variable than others. Contraction duration was the most consistent kinematic parameter among nectophores (Supplemental Figure 1), meaning that the jets produced by the nectophores within a species do not substantially change over time for normal swimming. However, kinematic parameters that quantify timing between adjacent nectophores (e.g., interpulse duration and offset time) and the time between successive pulses of any one nectophore (e.g., jetting frequency) are much less consistent.

None of the kinematic parameters differed significantly among the nectophores for each species (ANOVA, p > 0.05 for all species). Therefore, we averaged the kinematic parameters among all the nectophores of each individual to compare the parameters among species. Contraction durations were highly consistent within each species, however, they differed significantly between species (single factor ANOVA on log transformed data, p<0.05; Tukey-Kremer post hoc test, p < 0.05; Figure 3A). The contraction duration for *N. bijuga* from Friday Harbor was more than double the duration of *Nanomia* sp. from Panama. However, the water temperature in Friday Harbor was 21° C colder than Panama (8 ° C vs. 29 ° C). Based on this temperature difference we calculated a Q10 = 1.58 ± 0.09. To examine how contraction durations related to the nectophore size of the different species, we adjusted the rates of *Nanomia* sp. from Panama based on the calculated Q10 to 8° C (the ambient temperature of the other species). Contraction durations were strongly related to nectophore size (Regression P < 0.05, R^2^ = 0.93; Figure 3B). Interpulse duration also varied among species (Fig. 3C; single factor ANOVA on log transformed data, p<0.05), however the duration between pulses was not as tightly dependent on nectophore size as pulse duration (Fig. 3D). The jet frequency also varied among species (Fig. 3E; single factor ANOVA on log transformed data, p<0.05). While the two species with the largest nectophores had significantly lower frequencies (Tukey-Kremer post hoc test, p < 0.05), the regression with nectophore size among all the species was not significant (Regression analysis, p > 0.05).

**Fig. 3.**
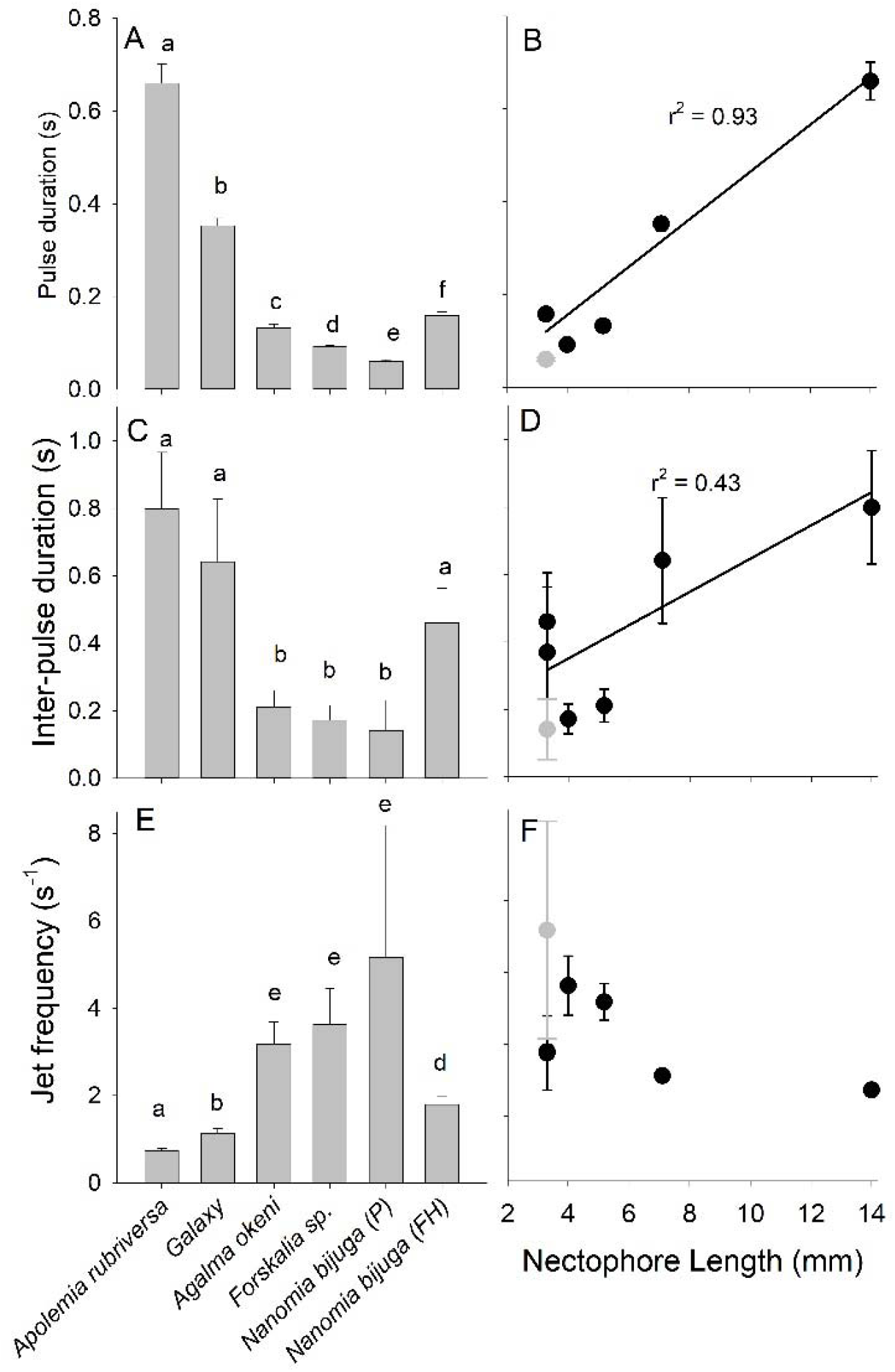
Nectophore kinematics among species (ordered from largest to smallest). Pulse duration of individual nectophore compared A) among species and B) versus nectophore length of each species. Interpulse duration of individual nectophores compared C) among species and D) versus nectophore length. Jetting frequency of individual nectophores E) among species and F) versus nectophore length. Grey circle is the nontemperature corrected rates for *Nanomia* sp. from Panama which was not included in the regressions. The temperature was adjusted to 8C for regressions. Error bars are st. dev.

To examine if adjacent nectophores on opposite sides of the nectosome coordinated their pulses, we compared the offset times between left and right nectophores for the Galaxy siphonophore, *Nanomia* sp. (Panama), and *A. rubriversa*; these were the only animals where we could resolve both sides of the nectosome (Fig 4). A negative vs. positive offset meant that the left nectophore pulsed before vs. after the right nectophore, respectively. The offset times between the left and right nectophores were highly inconsistent for all three species examined (Fig. 4). There was no consistent pattern for which nectophore pulsed first, with the left nectophores sometimes pulsing before the right, and also pulsing after the right just as often. In addition, the offset times were highly variable; occasionally the offset was equal to or greater than the interpulse duration of individual nectophores. Consequently, nectophores on opposing sides of the nectosomes do not appear to be coordinated. However, a comparison of the pulse frequency of nectophores on each side showed that the nectophores along opposing sides of the nectosome pulsed with the same frequency (Fig. 5). Therefore, the same amount of thrust would be produced on each side of the nectosome, enabling the siphonophore to maintain a straight trajectory.

**Fig. 4.**
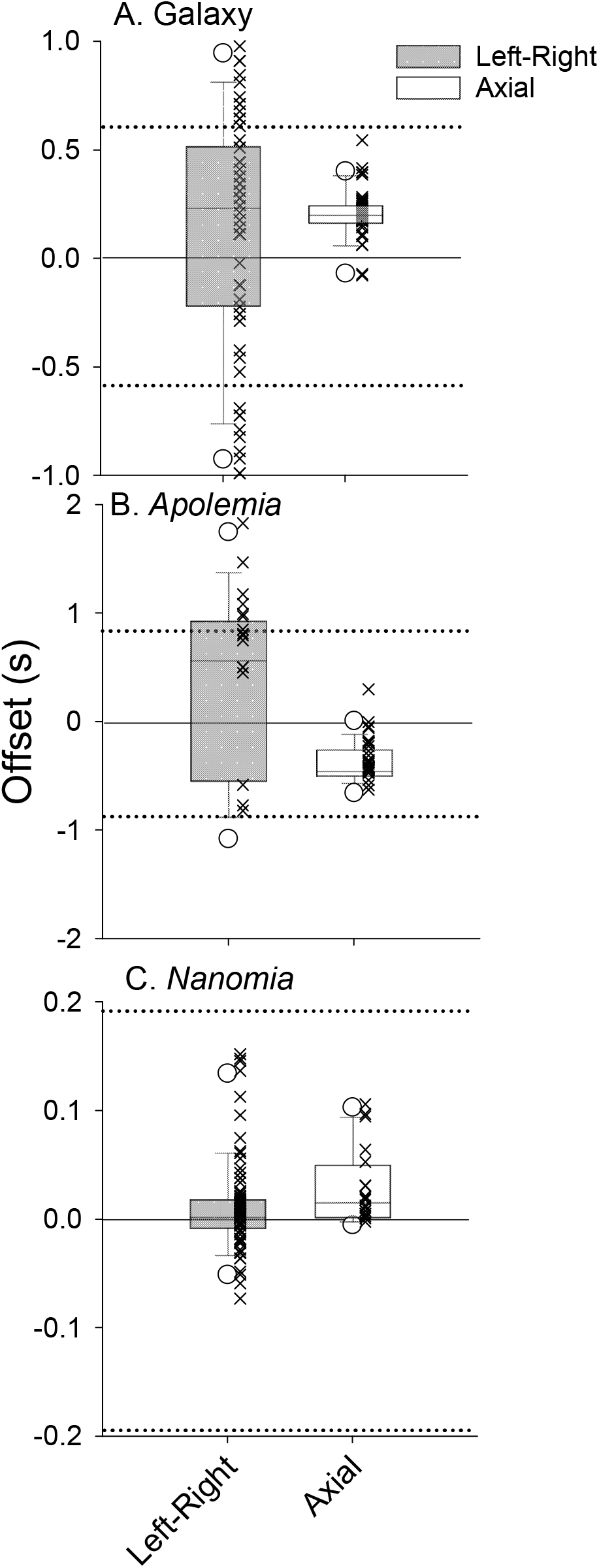
Box-plots of offset time between left and right nectophores and axially aligned nectophores for 3 species. X symbols are the individual data. Dashed line shows the interpulse duration. Negative offset means left (for Left-Right comparison) or leading (for Axial comparison) contracted before right or trailing nectophore, respectively.

**Fig. 5.**
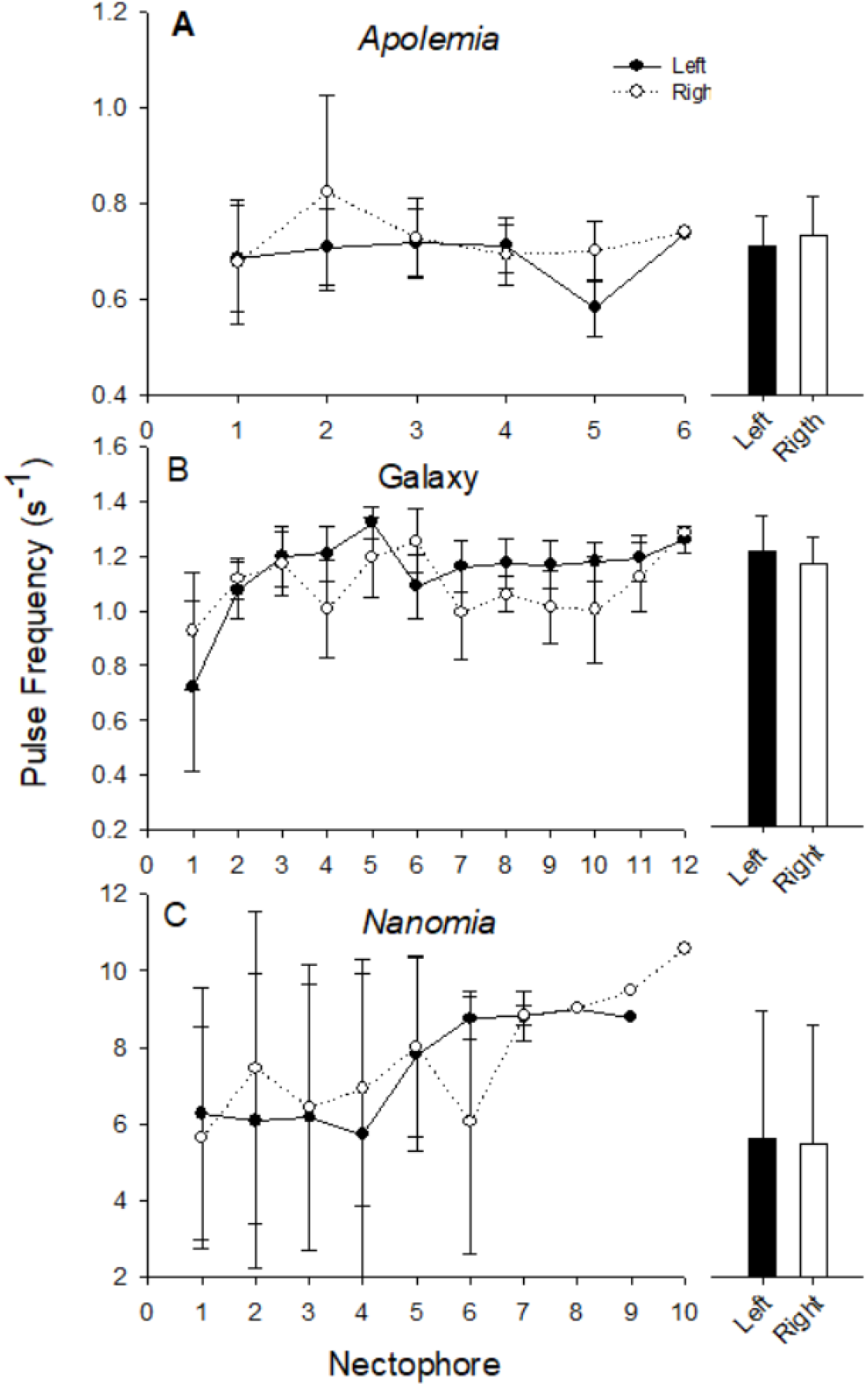
Mean pulse duration of nectophores along the left versus right side of the nectosomes. The closed vs. open symbols show the mean (± s.d.) of left vs. right nectophores, respectively, based on position averaged over replicate individuals. Bars show the mean (± s.d.) of all nectophores along the nectosome.

In contrast, nectophores aligned axially along the same side of the nectosome did appear to be consistently coordinated (Fig. 2, Fig. 4). To illustrate the coordination compared to the lack of coordination between the left and right nectophores, we similarly plotted the offset of adjacent nectophores along the same side of nectosome, and these plots show much less variable offset times. The leading nectophore always pulsed before the trailing nectosome (except for *A. rubriversa* which had the opposite sequence), and the offset was much shorter than the interpulse duration. The consistent offset times indicated that the nectophores were coordinated metachronally. A positive offset time meant that the metachronal wave traveled from the apex to the dorsal end of the nectosome (Fig. 4). The negative offset time of *A. rubriversa* indicated that the wave traveled from the dorsal end to the apex. The offset times varied among the species, and appeared to be related to the pulse duration of each species (ANOVA, p < 0.02, Holm-Sidak post-hoc comparison, p<0.05, Fig. 6A). The slope of the line of the offset time vs. the pulse duration was 0.45 indicating that the adjacent nectophores initiated their pulses about halfway through the pulse of the previous nectophore (Fig. 6B).

**Fig. 6.**
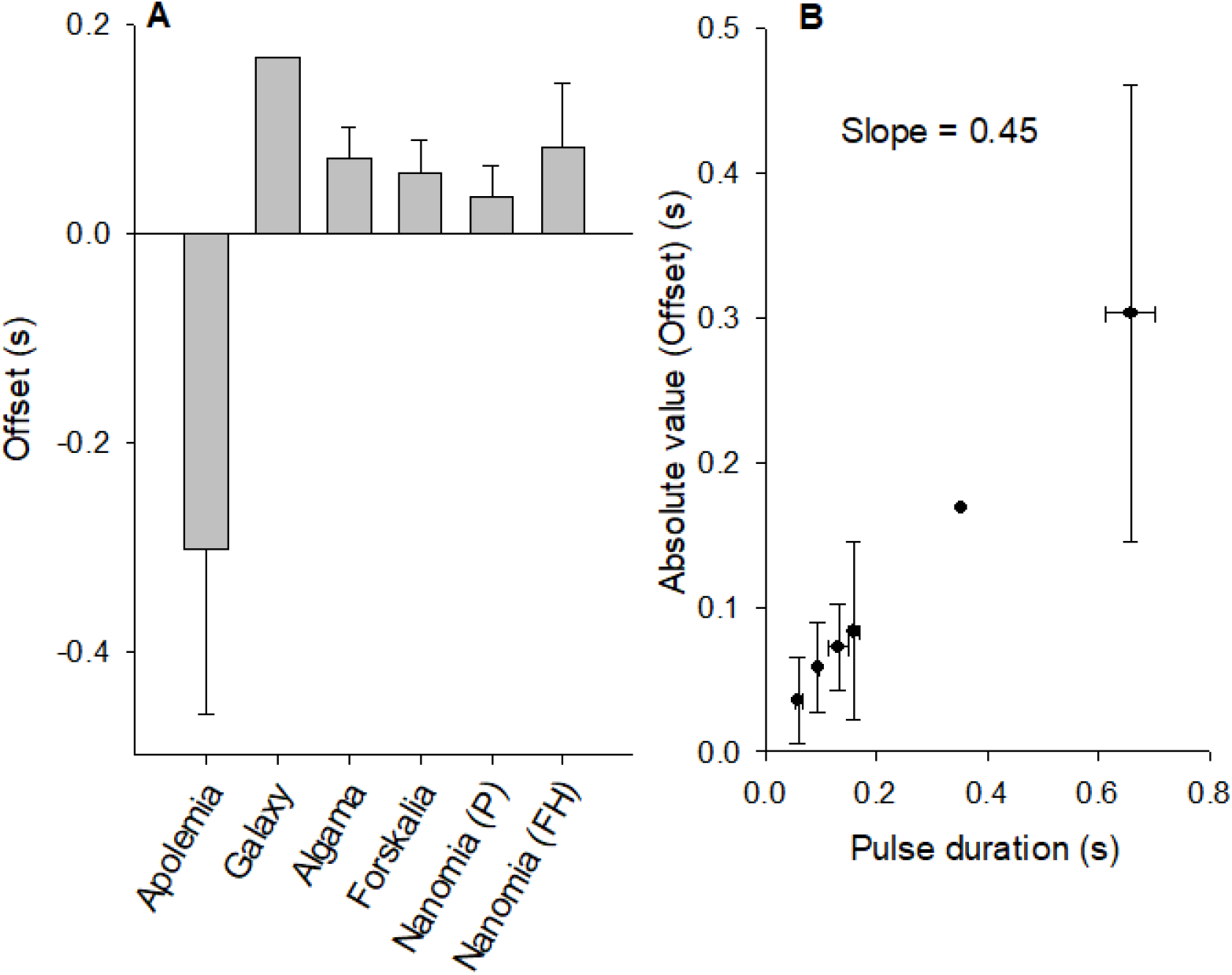
Comparison of offset between axially aligned nectophores (A among species and (B) versus pulse duration. Error bars show standa deviations.

Particle image velocimetry (PIV) of adjacent jets, where the trailing nectophore initiated its contraction halfway through the pulse of the leading nectophore, shows that the wakes of the two jets eventually interact downstream from the nectophore orifice (Fig. 7). Very few PIV sequences of asynchronous jets exist because in captivity, a siphonophore will exhibit mostly reactive escape swimming with synchronized jets. In their natural environment, as our data show, siphonophores use asynchronous jets during normal swimming. In Fig. 8 the siphonophore was starting from rest (v = 0) and its swimming speed was less than observed in the field. This would result in the jet wake being further apart and interacting less than during steady-state swimming. Despite this, we can see that the leading jet is fully developed when the trailing jet is initiated. Most of the flow of the leading jet was a well-organized linear jet moving in the same direction as the jet of the trailing nectophore.

**Fig. 7.**
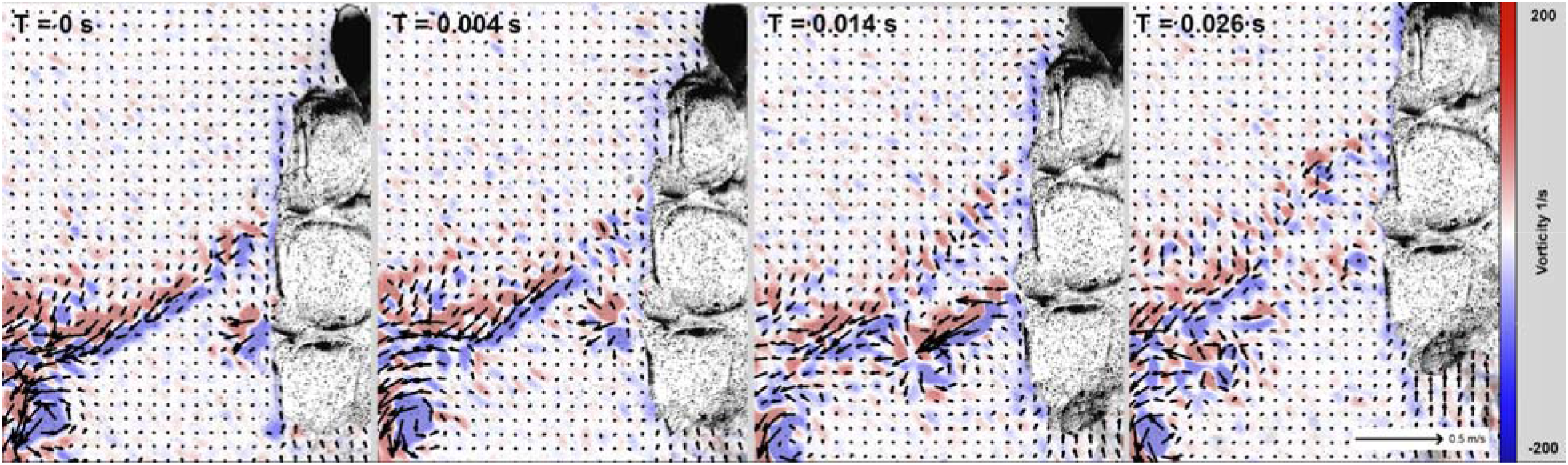
Sequential particle image velocimetry (PIV) of asynchronous jets of *Nanomia bijuga.*

## Discussion

Multi-jet swimming is a relatively rare propulsive strategy for animals, and is observed for only two known taxa: salps and siphonophores. Both taxa appear to synchronize their jets to escape predators (Mackie, 1964; Sutherland et al., 2017; Sutherland et al., 2019a), which maximizes thrust production and acceleration. However, for routine swimming that is often used for feeding and repositioning, the jets of both taxa are timed asynchronously (Costello et al., 2015; Sutherland et al., 2017). Asynchronous jetting has been shown to produce less thrust and lower swimming speeds, but is more efficient than synchronous jetting. Efficiency is thought to be enhanced because asynchronous jetting produces more steady swimming speeds (i.e.; fewer accelerations and decelerations), which serves to reduce drag (Du Clos et al., 2022; Sutherland et al., 2017). However, at the intermediate Reynolds numbers (Re) experienced by siphonophores and salps, other properties of the jets (e.g., timing and jet angle) can have important effects on drag and efficiency (Jiang et al., 2021). Previous studies have shown that the timing of most salp jets are uncoordinated (Sutherland et al., 2017), while siphonophore jets – mostly based on observations of *Nanomia bujiga* – appear to be coordinated (Du Clos et al., 2022). We show that for several physonect siphonophores, nectophores on the same side of the nectosome are well coordinated in a metachronal wave pattern. However, nectophores on opposing sides of the nectosome do not appear to be coordinated with each other. Metachronal waves are typically initiated near the apex and travel dorsally along the nectosome. *Apolemia* is an exception to this pattern because metachronal waves move from the nectosome base towards the apex. For all the species the nectophore pulses are not always tightly coordinated and some nectophores either miss a wave or pulse out of phase with their neighbors.

The advantages of metachronal coordination by nectophores remains unexplored. Wake interactions may influence efficiency or thrust production of the nectosome, however this has been rarely examined; among the few existing studies, most show that interactions between jets can reduce the thrust produced by a jet. For example, it’s been shown that both thrust and efficiency of jets decreased as two synchronized pulsed jets moved closer together and interacted more (Athanassiadis and Hart, 2016). In addition, it is known that the flow environment surrounding pulsed jets further affect the thrust production where parallel co-flow decreases thrust and counter-flow may increase thrust (Dabiri and Gharib, 2004; Krueger et al., 2003). Evidence from this study demonstrates that adjacent nectophores generally initiate their pulse halfway through the jet of the previous nectophore, and that the leading jet wake is linear in the co-flow direction (i.e.; same direction) of the adjacent nectophore when it initiates jetting (Fig. 6, T = 0.004 and 0.014). The wake of the second jet experiencing co-flow with an adjacent jet should therefore experience reduced thrust production (Dabiri and Gharib, 2004).

Orientation of jets, in addition to timing, plays an important role in thrust and efficiency (Jiang et al. 2021) because as jet angles increase (jets directed more laterally relative to the nectosome axis), their wakes interacted less. Jiang et al. (2021) found that two optima of jet angles exist depending upon the jetting parameters that are optimized. Larger jet angles (61-70°) optimized the quasipropulsive efficiency because these larger angles had fewer jet interactions and required less overall jet power to swim. Alternatively, smaller jet angles (34-45°) optimized Froude efficiency because jet interactions minimized wake kinetic energy, but required more jet power to overcome the drag associated with greater wake interactions. Interestingly, the wakes of many multi-jet salps appear to optimize quasipropulsive efficiency by directing their jets more laterally (Sutherland et al., 2017). This is intuitively appealing for salps that primarily swim to process fluid for feeding, and produce less powerful jets than siphonophores. In contrast, the sole role of siphonophore jets is to provide thrust for colony transportation. The jets of many siphonophores are angled more longitudinally with angles < 50°. These angles produce more wake interactions but result in higher Froude efficiencies. Therefore, the metachronal timing of the jets may serve to further optimize efficiency of jet interactions.

Contraction time (t) of individual nectophores also affects the thrust produced by each propulsive unit. The jet thrust emerging from a nectophore is determined by the mass and acceleration of the ejected fluid, and the nectophore size, orifice diameter, and contraction rate are the primary factors that affect the mass efflux from the nectophore (Colin and Costello, 2002; Daniel, 1983). We found that contraction times among the physonect species were directly related to nectophore size (x) (Fig. 3B). Longer contraction times serve to decrease acceleration, but larger sizes should also eject a greater volume of fluid. We can estimate the net effect of nectophore size on thrust (T) as (ρA_v_)(dV/dt)^2^ where ρ is the density of seawater, A_v_ is the area of the aperture opening, V is the volume inside the nectophore, and t is the contraction duration (Colin and Costello, 2002; Daniel, 1983). Therefore, if we assume that A_v_ scales with nectophore size and percent dV is constant among the species, we can estimate thrust value based on morphology of *N. bujiga.* Based on this, we found that thrust increased with nectophore size (x in mm) by the relationship T (μN) = 9.1x – 19.9 (with an r^2^ = 0.97). Therefore, despite the longer contraction times, thrust appears to increase with nectophore size by a factor of almost 10 due primarily to the greater fluid volumes ejected by larger nectophores. This indicates that both the number and size of nectophores comprising the nectosome will directly increase thrust produced by different siphonophores species. Note that this comparison would be illuminating with the other species we studied, however we lack the necessary morphological data to estimate the thrust as we did here.

Although nectophores along the same side of the nectosomal axis are generally metachronally coordinated, the degree of coordination can be inconsistent due to variability in contraction timing by adjacent nectophores. The loose coordination may be the result of the simple neuromuscular system of cnidarian siphonophores. The nectophores are attached to a central nectosome stem through which the lower nerve tract travels; the lower nerve tract then connects the nectophores and stimulates forward swimming. However, the speed and fidelity of neural signal transfer is limited for this comparatively simple neural system (Mackie, 1964; Norekian and Meech, 2020). Despite their neural simplicity, cnidarians such as siphonophores and medusae are able to swim effectively and their level of neuromuscular control appears adequate for the behavior requirements of swimming. In other words, while the coordination of nectophores is not perfect it may be “good enough” to achieve the behavioral requirements of siphonophore swimming. The limited behavioral observations of siphonophores have shown that they alternate between feeding bouts of motionless drifting with repositioning bouts requiring movement to a new location (Madin, 1988). The asynchronous kinematics we observed occurred during repositioning bouts. Even though the nectophores on opposite sides of the nectosome were not synchronized, the colonies were able to maintain relatively linear trajectories because the pulse frequencies of nectophores on both sides of the nectosome were equal. Coordination between sides along the nectosome does not appear necessary for achieving a linear heading. Likewise, the loose metachronal coordination of axial nectophores is adequate for effective repositioning of a siphonophore moving to a new feeding location.

Medusae demonstrate that, for individual cnidarians, propulsive mode is strongly related to foraging strategy (Costello et al., 2008; Dabiri et al., 2010). For individual medusae, ambushing species optimize thrust production over efficiency while feeding-current-foraging medusae optimize efficiency over thrust. We also believe that similar relationships may exist for siphonophores. Some siphonophores are known to migrate over long distances (Pages and Gili, 2014), while others are known to swim much more continuously than others (unpublished data). Nectophore design and operation are potential avenues for siphonophores to modulate tradeoffs between propulsive thrust and efficiency. Both thrust and efficiency (Jiang et al. 2021, DuClos et al. 2022) are affected by nectophore jet interactions, and our results clearly demonstrate the ability of siphonophores to coordinate these jet interactions. Although the complexities of coordinating multi-jet propulsion are substantial, comparison of multiple species is allowing us to establish an framework of how siphonophore colonies successfully accomplish this challenge.

## References

Athanassiadis, A. G. and Hart, D. P. (2016). Effects of multi-jet coupling on propulsive performance in underwater pulsed jets.

Colin, S. P. and Costello, J. H. (2002). Morphology, swimming performance and propulsive mode of six co-occurring hydromedusae. J. Exp. Biol. 205, 427–37.

Costello, J. H., Colin, S. P. S. P. and Dabiri, J. O. J. O. (2008). Medusan morphospace: phylogenetic constraints, biomechanical solutions, and ecological consequences. Invertebr. Biol. 127, 265–290.

Costello, J. H., Colin, S. P., Gemmell, B. J., Dabiri, J. O. and Sutherland, K. R. (2015). Multi-jet propulsion organized by clonal development in a colonial siphonophore. Nat. Commun. 6,.

Dabiri, J. O. and Gharib, M. (2004). Fluid entrainment by isolated vortex rings. J. Fluid Mech. 511, 311–331.

Dabiri, J. O. O., Colin, S. P. P., Katija, K. and Costello, J. H. H. (2010). A wake-based correlate of swimming performance and foraging behavior in seven co-occurring jellyfish species. J. Exp. Biol. 213, 1217–25.

Daniel, T. L. (1983). Mechanics and energetics of medusan jet propulsion. Can. J. Zool. 61, 1406–1420.

Du Clos, K. T., Gemmell, B. J., Colin, S. P., Costello, J. H., Dabiri, J. O. and Sutherland, K. R. (2022). Distributed propulsion enables fast and efficient swimming modes in physonect siphonophores. 119, 1–8.

Jiang, H., Costello, J. H. and Colin, S. P. (2021). Fluid dynamics and efficiency of colonial swimming via multijet propulsion at intermediate Reynolds numbers. Phys. Rev. Fluids 6,.

Katija, K., Roberts, P. L. D., Daniels, J., Lapides, A., Barnard, K., Risi, M., Ranaan, B. Y., Woodward, B. and Takahashi, J. (2021). Visual tracking of deepwater animals using machine learning-controlled robotic underwater vehicles. In 2021 IEEE Winter Conference on Applications of Computer Vision,.

Krueger, P. S., Dabiri, J. O. and Gharib, M. (2003). Vortex ring pinchoff in the presence of simultaneously initiated uniform background co-flow. Phys. Fluids 15,.

Mackie, G. O. (1964). Analysis of Locomotion in a Siphonophore Colony. Society 159, 366–391.

Madin, L. P. (1988). FEEDING BEHAVIOR OF TENTACULATE PREDATORS: IN SITU OBSERVATIONS AND A CONCEPTUAL MODEL L. p, Madin. Bull. Mar. Sci. 43, 413–429.

Norekian, T. P. and Meech, R. W. (2020). Structure and function of the nervous system in nectophores of the siphonophore Nanomia bijuga.

Pages, F. and Gili, J. (2014). Vertical distribution of epipelagic siphonophores at the confluence between Vertical distribution of epipelagic siphonophores at the confluence between Benguela waters and the Angola Current over 48 hours. Hydrobiologia 216, 355–362.

Pugh, P. R. (1975). The distribution of siphonophores in a transect across the North Atlantic Ocean at 32 N. J. Exp. Mar. Bio. Ecol. 20, 77–97.

Purcell, J. E. (1980). Influence of siphonophore behavior upon their natural diets: Evidence for aggressive mimicry. Science (80-.). 209, 1045–1046.

Purcell, J. E. (1981). Dietary composition and diel feeding patterns of epipelagic siphonophores. Mar. Biol. 65, 83–90.

Robison, B. H., Reisenbichler, K. R., Sherlock, R. E., Silguero, J. M. B. and Chavez, F. P. (1998). Seasonal abundance of the siphonophore, Nanomia bijuga, in Monterey Bay. Deep Sea Res. Part II Top. Stud. Oceanogr. 45, 1741–1751.

Sutherland, K. R., Weihs, D. and Sutherland, K. R. (2017). Hydrodynamic advantages of swimming by salp chains.

Sutherland, K. R., Gemmell, B. J., Colin, S. P. and Costello, J. H. (2019a). Propulsive design principles in a multi-jet siphonophore. J. Exp. Biol. 222, 1–8.

Sutherland, K. R., Gemmell, B. J., Colin, S. P. and Costello, J. H. (2019b). Maneuvering performance in the colonial siphonophore, Nanomia bijuga. Biomimetics 4,.

